# Megakaryocytes Display Innate Immune Cell Functions and Respond during Sepsis

**DOI:** 10.1101/742676

**Authors:** Galit H. Frydman, Felix Ellett, Julianne Jorgensen, Anika L. Marand, Lawrence Zukerberg, Martin Selig, Shannon Tessier, Keith H. K. Wong, David Olaleye, Charles R. Vanderburg, James G. Fox, Ronald G. Tompkins, Daniel Irimia

## Abstract

Megakaryocytes (MKs) are precursors to platelets, the second most abundant cells in the peripheral circulation. However, while platelets are known participate in immune responses and play significant roles during infections, the role of MKs within the immune system has not been explored. Here we utilize *in vitro* techniques to show that both cord blood-derived MKs (CB MKs) and MKs from a human megakaryoblastic leukemia cell line (Meg-01) chemotax towards pathogenic stimuli, phagocytose bacteria, and release chromatin webs in response to bacteria. Moreover, in patients with sepsis, we found that MK counts were significantly higher in the peripheral blood, and CD61^+^ staining was increased in the kidneys and lungs, correlated with the development of organ dysfunction. Overall, our study suggests that MK cells display basic innate immune cell functions and respond during infections and sepsis.

---

Megakaryocytes (MKs) are commonly recognized as key participants in hemostatic processes through the production of platelets^1–2^. In addition to their presence in the bone marrow, MKs can also be located in the lungs, lymph nodes, spleen, and liver during extra medullary hematopoiesis^3–7^. MKs have also been reported to be significantly increased in the lungs during severe pulmonary inflammation, such as acute respiratory distress syndrome (ARDS), when they are believed to promote inflammation via the release of platelets^8–10^. The current paradigm by which MKs are increased in the lungs during ARDS revolves around MKs passive escape from the bone marrow, entrance to arterial circulation, and passive mechanical entrapment within the microcirculatory bed of the alveoli^4,11–12^.

The participation of MKs in immune responses is suggested by several anecdotal observations^19^. Maturing MKs express both major histocompatibility complex (MHC) class I and II molecules and a variety of toll-like receptors (TLRs) on their cell surface ^13–19^. MKs, just like platelets, also contain various granule types, including lysosomes, which participate in the endocytosis and degradation of pathogens^20^. MKs can play antigen-presenting-cell (APC) roles and stimulate Th-17 responses in lupus^13,21^. Thrombocytes, the amphibian equivalent of the mammalian MK/platelet, actively phagocytose live bacteria^23–25^. In mammals, MKs internalize viruses, including dengue virus and HIV, and multiple case reports show evidence for MKs containing fungi^26–29^. Although such reports provide sporadic support for an active role for MKs in the immune responses, this function of MKs has not been tested systematically.

Here, we show that the human MKs can engulf pathogens, release chromatin nets, and undergo chemotaxis in gradients of standard chemoattractants. Moreover, we find that during sepsis, the number of large CD61^+^CD41^+^ cells increases in peripheral circulation, and the number of large CD61^+^ cells increases in peripheral organs. These numbers are higher during acute kidney injury (AKI), ARDS, and in disseminated intravascular coagulation (DIC).

## RESULTS

### MKs are phagocytic

We tested the capacity of CB MKs and Meg-01 cells to engulf *E. coli, S. aureus,* or *S. pyogenes* (**Figure 1**). We used light microscopy to verify the association of bacteria with the cell membrane (**Figure 1A**) and transmission electron microscopy to confirm the internalization of bacteria by MK and platelets (**Figure 1B**). Labeling the pathogens with pHrodo, a marker that fluoresces red following acidification, confirmed the uptake of the bacteria within phagosomes (**Figure 1C** and **Figure S2**). Although the Meg-01 cells are noted to have strong auto fluorescence (both red and green), the observation of the rod-shaped, bright red stained *E. coli*, further confirms the internalization of the bacteria by the Meg-01 cells. Interestingly, electron micrographs of a spontaneously contaminated CB MK culture at day 14 of differentiation revealed one large cell with multiple bacteria of unknown origin within an expansive vacuole (**Figure 1B**).

**Figure 1.**
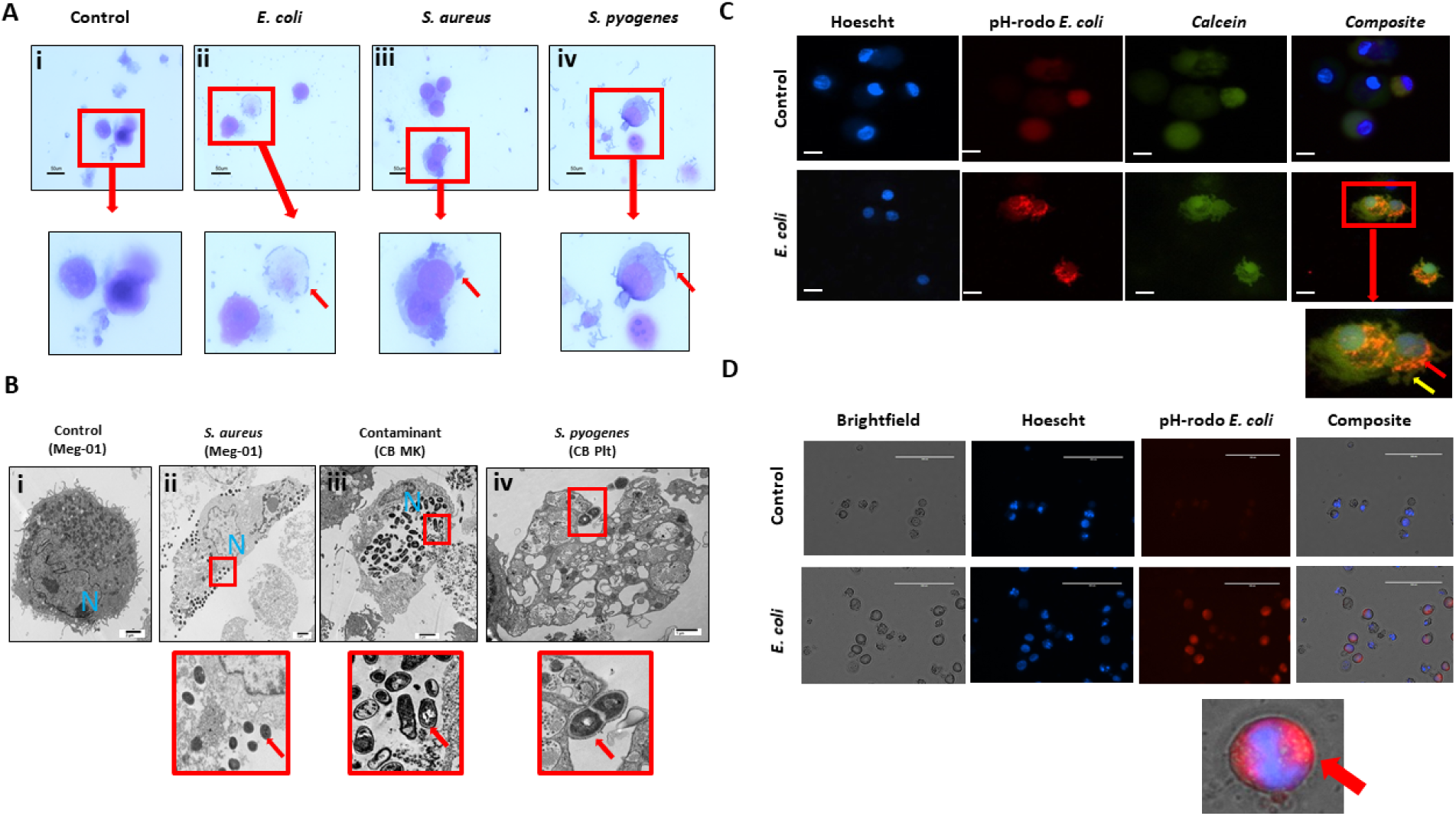
MKs are capable of phagocytosis of pathogens. (A) Meg-01 cells were co-incubated with E. coli, S. aureus, and S. pyogenes. Light microscopy with diff-quick staining shows that bacteria are associated with the cytoplasm and cell membrane of the cells. (B) Transmission electron microscopy of both Meg-01 and CB MKs exhibiting bacterial association with the cell membrane as well as internalization into vacuoles within the cytoplasm. Panel i is a control Meg-01 cell, while panel ii is a Meg-01 cell that was co-incubated with live *S. aureus.* Panel ii shows a CB MK with multiple bacteria within a large cytoplasmic vacuole from a contaminated cell culture. Panel iii is from a CB MK culture co-incubated with live *S. pyogenes* showing association of the bacteria with a platelet cell membrane. (C) CB MKs were co-incubated with live pHrodo-conjugated bacteria. This is a representative image of a control cell along with a cell co-incubated with *E. coli*. (D) Meg-01 cells were co-incubated with pHrodo-conjugated live *E. coli* and then imaged. Bacteria, red arrow; pseudopodes, yellow arrow.

We incubated CB MKs at different stages of differentiation (day 0-14) with bacteria. We confirmed appropriate MK maturation by measuring surface-marker expression using flow cytometry (**Figure S3**). We found that CB MKs were capable of phagocytosis of *E coli* starting at day 10 of differentiation (**Figure S2A**). This corresponds temporally to the onset of P-selectin glycoprotein ligand 1 (CD162) and MHC Class II (HLADR) expression (**Figure S3**). Cells also appeared to be capable of internalizing zymosan particles during overnight incubation, although this response was less robust compared to that against bacteria (**Figure S2B**). Meg-01 also phagocytose bacteria. Interestingly, Meg-01 phagocytose *beta-hemolytic E. coli* more efficiently compared to *non-beta-hemolytic E. coli* (data not shown), suggesting that various specific receptors and pathogen-cell interactions take place prior to internalization.

### MKs undergo chemotaxis

We tested the ability of Meg-01 cells to chemotax towards LPS and zymosan particles in in microfluidic as well as traditional transwell assays (**Figure 2**). In microfluidic assays, we observed Meg-01 chemotaxis at single cell resolution and distinguished three phenotypic groups. Cells in the first group migrated through the side channels and entered the circular reservoirs. Cells in the second groups remained in the side channels and extended projections into the side channels. Finally, cells in the third group remained in the side channels (**Figure 2B-C** and **Figure S5**). Strikingly, the size of the cells was not an impediment for cell migration and large Meg-01 cells, up to 75 μm in diameter, actively migrated through 4.5 μm high and 10.5 μm wide channels. We observed nuclei and organelle displacement (**Video 1-3**), with some moving cells carrying zymosan particles within them (**Video 4**). A small proportion of cells did not fully traverse the channel, but instead extended portions into side channels containing chemoattractant. These projections released small cell fragments (platelets or apoptotic bodies) towards the stimulus (**Figure 2D** and **Video 5**). We also observed cells migrating into the side channels and then releasing platelets or vesicles, effectively obstructing the side channel and preventing other cells from entering (**Video 6**).

**Figure 2.**
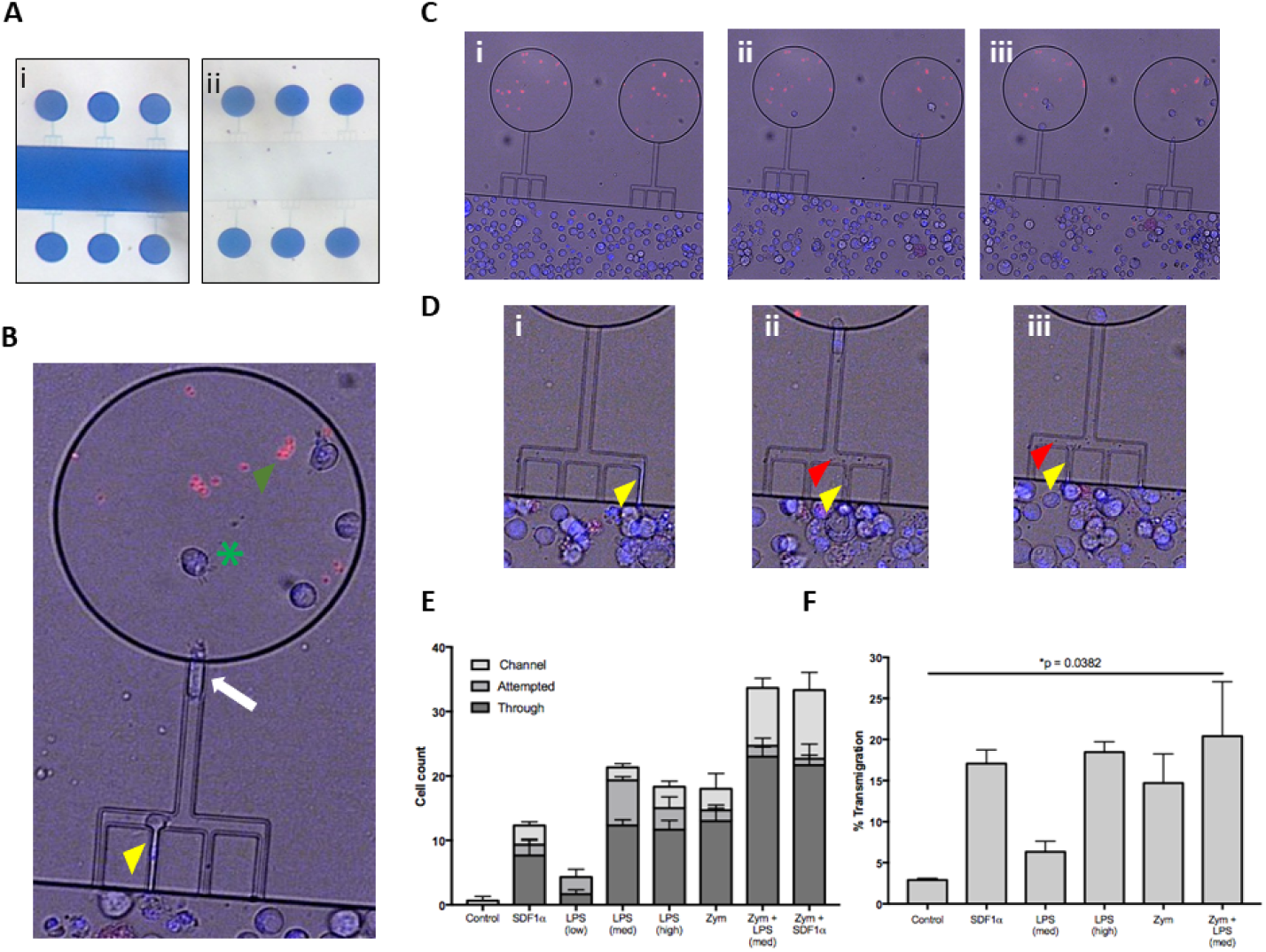
Meg-01 cells are capable of chemotaxis to pathogenic stimulus. Meg-01 cells were tested for their ability to chemotax towards LPS and zymosan particles. (A) A microfluidic device was used for part of the chemotaxis experiments. In this device, the main channel is connected to a circular reservoir by four 6 um channels and a larger 8 um connecting channel in a comb-like arrangement. The device is first primed with the condition (panel i) and the main channel is then flushed with media in order to create a concentration gradient from the ‘lollipops’ into the main channel (panel ii). (B) The MKs are stained with Hoechst for positive identification and then manually tracked. The behavior of the MK was divided into 3 categories: cells attempting to enter the channel (yellow arrowhead), cells inside the channel (white arrow), and cells through the channel and inside the reservoir (green arrowhead). Zymosan particles are marked with a green asterisk. (C) Time lapse image of MKs migrating into the lollipops which are primed with LPS (360 pg/mL) and zymosan particles. (D) Close-up of time-lapse image were MKs are observed to attempt to enter the channel, extend a portion of the cell into the side channel (yellow arrowhead), and then bud off small platelet-like particles (red arrowhead). (E) Bar graph representing MK chemotaxis within the microfluidic device. (F) Bar graph representing MK chemotaxis within a transwell device, confirming the same observation as within the microfluidic device. LPS low, 22 pg/mL; LPS med, 220 pg/mL; LPS high, 2.2 ng/mL; Zym, zymosan particles; Zym+LPS, zymosan particles with 220 pg/mL LPS. Bar graphs are the mean with standard error bars.

Chemotaxis of Meg-01 and MK cells towards the various stimuli in the microfluidic assay was consistent with traditional transwell assays (**Figure 2E-F**). In transwell experiments, the fraction of Meg-01 cells that migrated towards LPS at concentrations of 220 pg/mL and 2.2 ng/mL was 4.1-8.5 % and 16.0-21.0 %, respectively. The fraction of cells migrating towards zymosan particles was comparable: 7.6-21.8%. When LPS or SDF1-α was combined with zymosan particles, the average chemotaxis fraction increased slightly (7.1-33.7%). The positive control chemotaxis response of Meg-01 cells towards SDF1-α was between 13.7-20.4 %, consistent with previous reports.^35^

### MKs release chromatin webs

We observed that Meg-01 cells incubated with live bacteria or LPS change their cell morphology and release histone-decorated chromatin webs (**Figure 3**). Measurements of extracellular double stranded-DNA (dsDNA) in the supernatant show proportional increase of chromatin webs release with the concentration of LPS (**Figure 3B**). Fluorescent imaging of CB MKs co-incubated with live pHrodo conjugated *E. coli* helped visualize the chromatin webs and their filamentous structure (**Figure 3C**). During the early phase of chromatin web release, the nucleus of the cells is also stained by the Hoechst dye, which disappears in the late stages, consistent with recent reports in the context of neutrophil extracellular trap (NETs) release^36^. Co-incubation with heat-killed *E. coli* resulted in a less robust response compared to live bacteria (data not shown), consistent with literature on NET formation being dependent on bacterial motility^37^. Immunofluorescent imaging confirms the presence of extracellular histones and myeloperoxidase along with the chromatin webs (**Figure S6**).

**Figure 3.**
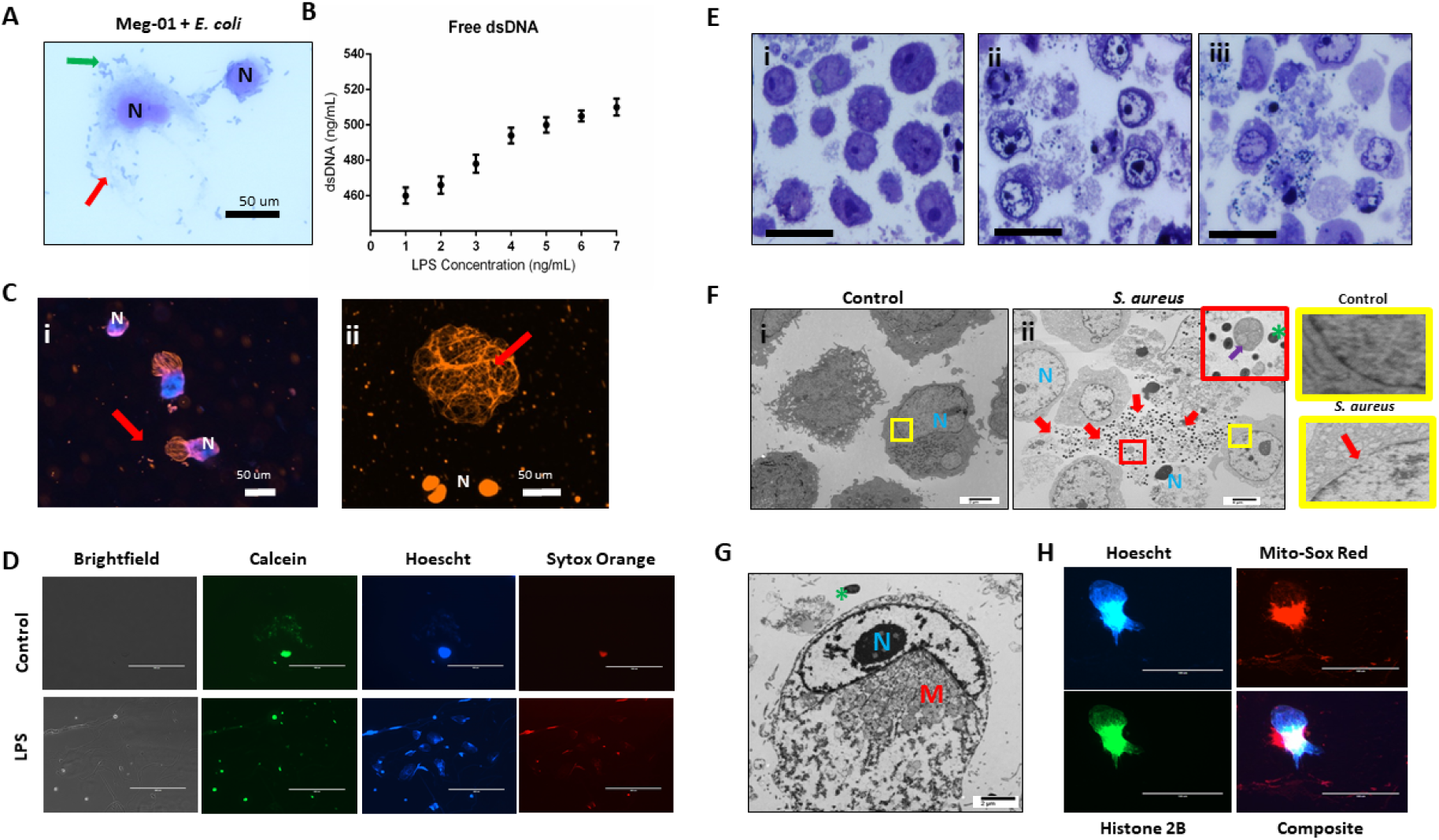
MK release chromatin webs. CB MKs and Meg-01 cells were observed to release chromatin webs in response to pathogenic stimulus. (A) Meg-01 cells co-incubated with live *E. coli* and undergo cell lysis (diff-quick stain). Green arrow: bacteria; red arrow: extracellular cytoplasm. (B) The amount of chromatin released from Meg-01 cells after incubation with increasing concentrations of LPS was quantified using a PicoGreen assay. (C) CB MKs release chromatin webs after incubation with live pHrodo-conjugated *E. coli* produce. Live cells are stained blue with Hoechst dye. Chromatin webs are stained orange with Sytox dye. Subpanel i depicts two CB MKS with nucleus (blue) and chromatin webs (orange). Subpanel ii depicts 4 CB MKs, one that has released a chromatin web (top) and three dead cells with stained nuclei (bottom). (D) Meg-01 cells in media have intact cell membranes and proplatelet buddings (calcein staining). Cells incubated with LPS have scant calcein and abundant sytox (chromatin) staining. (E) Meg-01 cells incubated with various live bacteria display swollen nuclei, broken nuclear membranes, chromatin webs, extracellular granules, and bacteria associated with intra- and extracellular contents (light microscopy). Panel i is the control, and panels ii and iii are co-incubated with *E. coli* and *S. aureus*, respectively. (F) Transmission electron microscopy (TEM) of Meg-01 cells in media (control) reveal an intact nuclear membrane (panel i) and limited extracellular content. Meg-01 cells co-incubated with live bacteria display swollen nuclei (N), broken nuclear membranes, extracellular cytoplasmic contents (including granules and mitochondria), and an abundance of bacteria primarily associated with this extracellular content (red arrows – panel ii). The red magnified section in panel ii demonstrates the presence of extracellular mitochondria (purple arrow) and the yellow magnified sections on the right demonstrate an intact nuclear membrane in a control cell (top right) and a cell co-incubated with bacteria that has a break in the nuclear membrane (bottom right). (G) TEM image of a Meg-01 cell co-incubated with live *E. coli* exhibiting a swollen nucleus and a rearrangement of mitochondria surrounding the nucleus. (H) Meg-01 cells transfected with Bacmam H2b-GFP released chromatin webs that were both positive for DNA (Hoechst) and histone 2B. Mitochondrial staining with MitoSox red shows active mitochondria in a perinuclear arrangement, confirming the TEM findings from panel G.

Electron micrograph imaging of Meg-01 cells co-incubated with live bacteria confirmed the presence of bacteria entangled within the chromatin webs along with various organelles, including extracellular mitochondria, granules, and nucleosomes (**Figure 3E-F**). Calcein staining demonstrates the lack of cell membrane around the chromatin webs and helps differentiate between cell death with intracellular content release. Calcein staining also differentiates from pro-platelet budding events, when the platelet buds are stained by calcein and are negative for DNA (**Figure 3D**). Other cell morphology changes consistent with both cell lysis and extracellular trap formation included swollen nuclei, breakdown of the nuclear membrane, decondensed chromatin, and re-localization of the cytoplasmic organelles and mitochondria ^38,39^. Furthermore, Meg-01 cells expressing GFP-H2B were imaged actively releasing their intracellular contents; which confirmed the perinuclear rearrangement of organelles (notably, mitochondria) (**Figure 3G-H**). In the case of MKs incubated with LPS, chromatin webs were released along the border of a hydrophobic pen marking on a glass slide. This was consistent with previous publications reporting NET formation upon contact with hydrophobic surface materials^40^.

### MKs are increased in the peripheral venous circulation during sepsis

We employed imaging flow cytometry to probe the presence of MKs in peripheral blood samples from patients. Circulating MKs were on average 10 µm in size and defined by the presence of CD61 and CD41 markers and Draq5 staining of the nucleus. We differentiated MKs from platelet-leukocyte aggregates by the distribution of markers, which was uniform throughout the cell membrane for MK cells and punctate when a platelet attaches to a lymphocyte (**Figure 4A**). The number of CD61^+^CD41^+^Draq5^+^ cells were significantly higher in peripheral circulation in patients with sepsis compared to controls (p = 0.05; 9565 <u>±</u> 1675 MK/mL vs 3502 <u>±</u> 741 MK/mL, respectively) (**Figure 4Bi**). Interestingly, MKs appeared to specifically be significantly increased in patients with ‘complicated’ (n = 9, including 2 follow-up counts on complicated patients) versus ‘uncomplicated’ sepsis (n = 4) (p = 0.01; 11077 <u>±</u> 6256 MK/mL vs 3587 <u>±</u> 1219 MK/mL, respectively) (**Figure 4Bii**), suggesting that there may be a correlation between the number of MKs in the peripheral venous circulation and the development of AKI or ARDS. When further subdividing the sepsis patient population by the source of the infection, patients with gram negative bacterial infections (n = 3) had significantly higher MK counts compared to those with gram positive bacterial infections (n = 7) (p < 0.01) but not those with the mixed infections (n = 3) (p = 0.13) (15070 <u>±</u> 5158 MK/mL, 5140 <u>±</u> 2082 MK/mL, 10146 <u>±</u> 7675 MK/mL, respectively) (**Figure 4Biii**).

**Figure 4.**
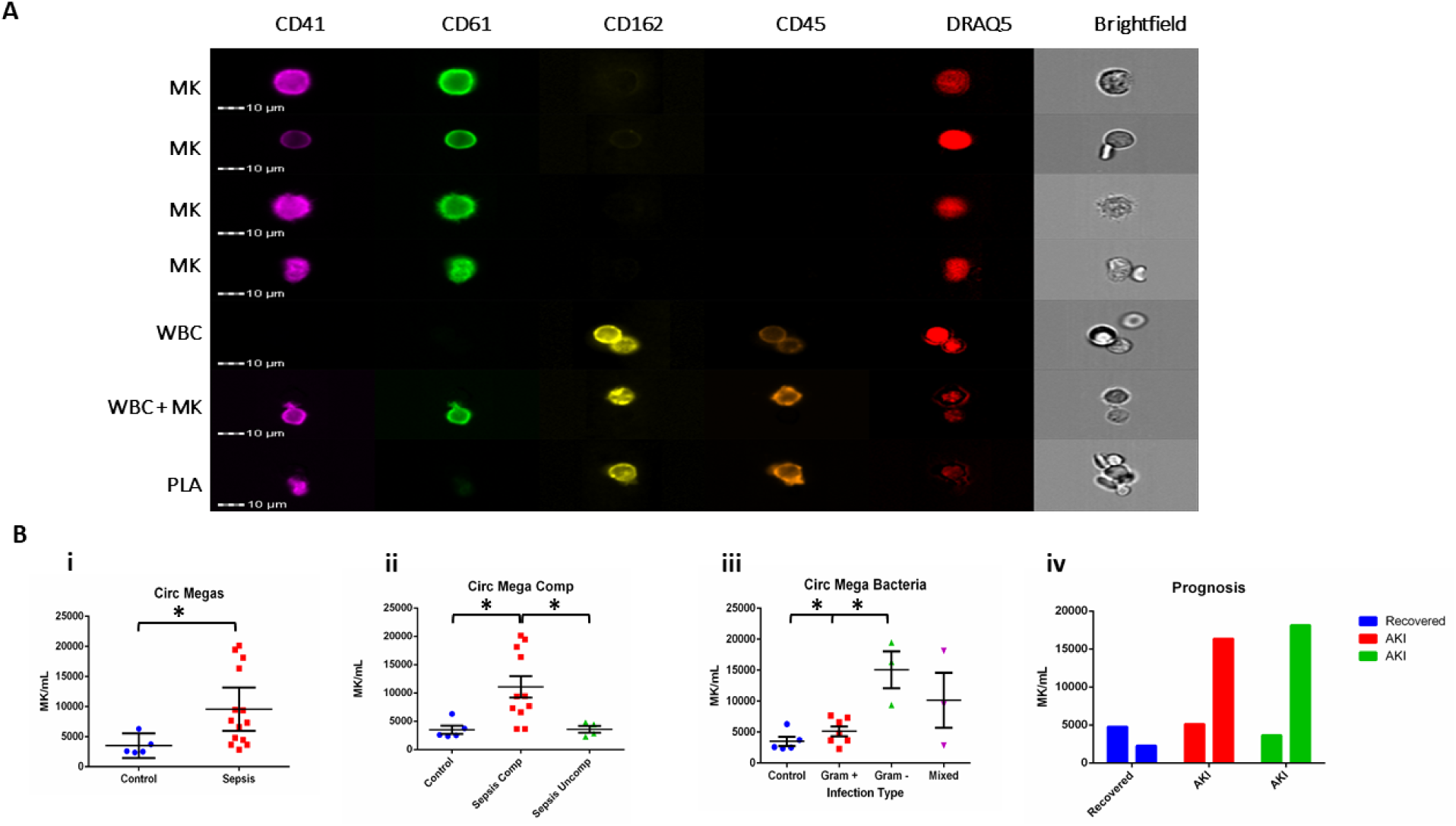
MKs are present in increased amounts in the circulation during sepsis. Samples from patients diagnosed with sepsis were evaluated for the presence of possible MKs. MKs were identified in the peripheral circulation based on the simultaneous expression of CD41, CD61, and DRAQ5. (A) Imaging flow cytometry was used to identify and quantify circulating CD41+CD61+Draq5+ cells in the peripheral circulation. CD45 and CD162 were used as white blood cell markers. The top 4 panels show examples of MKs, while the bottom three panels show examples of two white blood cells, a white blood cell attached to a MK, and a white blood cell attached to a platelet (PLA). The cells that stain negative for all stains in the images are most likely red blood cells.). (B) Circulating MKs were significantly higher in the peripheral circulation in patients with sepsis (i). The amount of MKs was correlated with sepsis-related complications, including ARDS and AKI, with ‘complicated’ sepsis having significantly higher MKs than ‘uncomplicated’ sepsis (ii). MKs were also noted to be higher in gram negative and mixed infections, as compared to gram positive only infections (iii). Sequential blood samples on day 1 and 3 of sepsis hospitalization suggested that there also may be a correlation with recovery and the development of sepsis complications, such as AKI (iv). Significance is calculated via student t-test with significance being defined as p < 0.05 (*). No statistical analysis was performed on B(iv) due to the small sample size (n = 1 per group).

### Automated analyzers fail to identify circulating MKs

We tested whether standard automated blood counting methods could identify circulating MKs (**Figure S1**). In an automated CBC analyzer, pure CB MKs were detected and categorized as an unknown type of white blood cells (WBCs) with an error notification. However, when CB MKs were spiked into whole blood samples, the automated analyzer categorized the additional cells as neutrophils, measured a corresponding increase in the WBC numbers, and did not display an error message (**Figure S1D**). Meg-01 cells could not be counted or analyzed accurately by the automated analyzer, neither as a pure population nor when spiked into whole blood. This is likely due to the differences in size between these two populations. While CB MKs are 10-30 µm and resemble small, granular lymphocytes on light microscopy, Meg-01 cells range from 15-75 µm and are often found in large clusters.

### Large numbers of circulating MKs correlate with worse prognosis in sepsis patients

To investigate whether MK enumeration in the circulation might be used to predict prognosis, we analyzed samples from three sepsis patients at 2 time-points during their hospital stay. We observed a decrease in circulating MKs in the patient that recovered and was discharged from the hospital, as compared to the two patients that remained in the hospital for multiple weeks and developed AKI (**Figure 4Biv**). Platelet count did not significantly differ in sepsis versus control patients (p = 0.98; 251.1 <u>±</u> 207.6 plt x 10^6/mL and 253.6 <u>±</u> 34.5 plt x 10^6/mL, respectively), whereas there was a significant difference between sepsis and control patients in total white blood cell counts (p = 0.015; 14.7 <u>±</u> 9.6 WBC x 10^6/mL and 5.2 <u>±</u> 1.1 WBC x 10^6/mL, respectively). There was no correlation between circulating MKs and platelet or white blood cell count (**Figure S7**). Interestingly, the circulating CD61^+^CD41^+^Draq5^+^ cells identified in the patient samples were CD162^−^, which may indicate that adult MKs have a different surface phenotype to neonatal MKs, which are CD162^+^ (**Figure S2**). Preliminary experiments aimed at identifying circulating MKs in neonatal intensive care unit (NICU) patients revealed that cells from these children appeared to be both CD34^+^ and CD162^+^ in addition to being CD41^+^Draq5^+^ (data not shown), suggesting that circulating MK phenotype may vary with the age of the patient. While the sample size in this exploratory study is too small to draw definitive conclusions, these data suggest that circulating MKs should be further explored in the context of sepsis-related complications.

### Platelets and MKs are present in peripheral organs during sepsis

To investigate whether CD61^+^ cells were increased in the peripheral organs of patients with sepsis, we performed detailed histological analysis of autopsy samples from sepsis patients and non-sepsis controls. CD61^+^ cells were defined as MKs when they were large with multi-lobular dark staining nuclei (as seen in **Figure 5A** iva-b). Per 40x field of view (FOV) of lung sections, large CD61^+^ MKs were significantly increased in the alveoli of septic patients (n = 8) as compared to controls (n = 4) (p = 0.01: 0.5 <u>±</u> 0.1 MK/FOV vs 1.4 <u>±</u> 0.2 MK/FOV, respectively) (**Figure 5A-B**). Comparing H&E stained sections to corresponding CD61 IHC image showed that although MKs could generally be identified on H&E by their large, darkly-stained nucleus, this was not always the case. This demonstrates the importance of utilizing IHC for accurate cell-type assessment in these types of analysis (**Figure 5A**).

**Figure 5.**
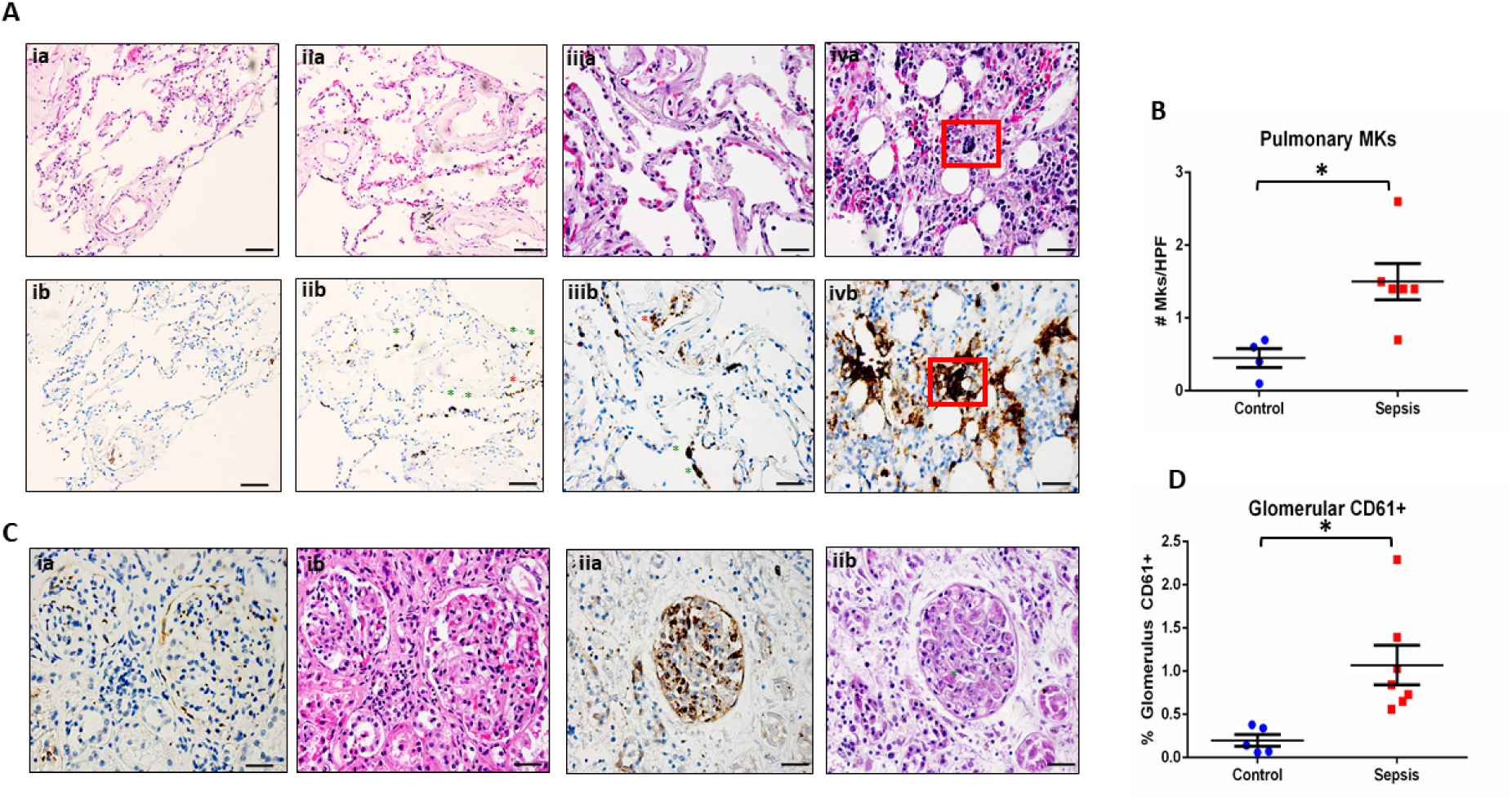
MKs are present in increased amounts in the peripheral organs during sepsis. Autopsy pathology samples from patients that died from sepsis were evaluated for MKs. (A) MKs in the lungs were defined as being large cells with dark, homogenous CD61+ staining (brown). An example of a lung image from a patient that died from heart disease as a negative control is shown in panel ia (hematoxylin and eosin, H&E) and ib (CD61). In these images there are no MKs observed. A representative image of two lung images from a patient that died from sepsis are shown in panel iia/iiia (H&E) and iib/iiib (CD61), showing multiple MKs (green asterisks). Platelet staining is also noted (red asterisks). While in some cases, a large dark staining basophilic nucleus can be seen in the H&E in the area of the CD61 staining, this is not always the case and may be due to sectioning through the sample and imaging different planes. This also suggests that CD61 may be more accurate when counting MKs than H&E. A sample of bone marrow from a sepsis patient is shown in panel iva (H&E) and ivb (CD61) as an example of a MK with CD61 staining as a positive control (red box). (B) Glomeruli were also evaluated for the presence of increased CD61 staining. Individual MKs could not be reliable counted within the MKs, therefore percent of CD61 stain/glomeruli was evaluated. A glomerulus from a control patient is shown in panel ia (CD61) and ib (H&E), with minimal CD61 staining. This is in contrast to a glomerulus from a patient with sepsis and disseminated intravascular coagulation (DIC), which has much higher amounts of CD61 staining along with microvascular thrombi within the glomerular capillaries (green asterisk). Upon evaluation, there was significantly higher pulmonary MKs (C) and glomerular CD61 staining (D) in the sepsis patients as compared to control. Significance is calculated via student t-test with significance being defined as p < 0.05 (*). Scale bars are: Ai-ii, 100 μm, Aiii-iv and C, 50 μm.

One ‘control’ patient (ID C2) was removed from pulmonary MK analysis because, although the cause of death was cardiac failure, a patchy bronchopneumonia was diagnosed at autopsy; this patient also had an increased number of MKs in the lungs (5.4 ± 3.5 MK/FOV), suggesting that MKs may be increased within the pulmonary parenchyma during localized as well as systemic infections. One patient (ID 4) that died from sepsis complicated by DIC was noted to have severe pulmonary inflammation and hemorrhage, as well as a micro-abscess, as defined by an increased number of neutrophils and macrophages in the lung (**Figure S8**). Gram staining and CD61 IHC identified intracellular gram-positive cocci, along with multiple large, homogeneously stained, CD61^+^ cells concentrated in the center of the abscess. This suggests that platelets, and possibly MKs, may have actively migrated out of the intravascular space and into the center of the abscess in response to inflammatory or bacterial signals.

Scoring of kidney sections revealed that renal glomeruli from septic patients (n = 7) also had a significantly increased amount of CD61+ positive staining than the controls (n = 5) (p = 0.018; 0.16 <u>±</u> 0.08% CD61/Glomerulus vs 1.07 <u>±</u> 0.23% CD61/Glomerulus, respectively) (**Figure 5C-D**). Comparing matched H&E and CD61 IHC glomeruli staining, it was not possible to differentiate with certainty whether large CD61+ areas corresponded to MKs or whether they represented platelet-rich fibrin thrombi (**Figure 5C**). One patient, who exhibited substantial microvascular thrombosis within the kidney and was diagnosed with disseminated intravascular coagulation (DIC), had significantly elevated CD61 staining (2.29 <u>±</u> 3.0 % CD61/Glomerulus), consistent with the presence of platelet-rich micro thrombi within the glomerular capillaries (**Figure 5D**).

## DISCUSSION

We found that MK cells display several immune cell functions, including chemotaxis, phagocytosis, and the release of histone-decorated chromatin webs, besides the traditional role in the production of platelets^2^. We demonstrate that, not only do MKs internalize bacteria, but they also localize the bacteria to acidified, lysosomal-type granules, which is indicative of an active killing process^44,45^. Our finding complements earlier observations that MKs contain various intracellular pathogens, including dengue virus, human immunodeficiency virus, and aspergillus^26–29^. The MK immune function complements earlier reports for platelets being ‘first responders’ to microbes and inflammatory insults, stimulating the recruitment and activation of white blood cells, and even directly trapping pathogens themselves ^41–43^. We also found that that Meg-01 cells undergo active chemotaxis towards common inflammatory stimuli^35^ and can migrate through channels with cross section smaller than 50 μm^2^. This size is comparable to that of small capillary vessels our observation raises the possibility that MKs are not ‘trapped’ in microcirculatory beds, but instead they may move through and diapedese into surrounding tissues where they participate in inflammation and regeneration^3,47–51^. Additionally, the observed directional budding of platelets into the channels containing the chemotactic agent and “obstruction” of the channels by MKs and platelets and/or vesicles, suggest an ability to contribute to decreased blood flow within small capillary vessels^10,43,52–55^.

We also demonstrate that MKs can release intracellular contents and generate chromatin webs. This feature is shared with an increasing number of innate immune cells that have been demonstrated to release their chromatin decondensed, including neutrophils, eosinophils^39,55^, and monocytes/macrophages. While the neutrophil-derived extracellular traps (NETs) have been shown to have antimicrobial function, the release of chromatin webs from MKs may be an essential part of the immune and coagulation systems. It is also possible that MK-derived chromatin webs participate in disease processes, including ARDs and sepsis, when there are increased numbers of MKs within the pulmonary parenchyma^53,66–67^.

The participation of MKs in immune responses is important in the context of our finding of large numbers of MKs in the peripheral circulation and peripheral organs in patients with sepsis. These findings are consistent with previous reports showing that MKs numbers are increased in the peripheral circulation in neonates with sepsis, in the renal glomeruli of adults with sepsis, and in the lungs in patients with sepsis and acute respiratory distress syndrome (ARDS)^70–71^. While the clinical data presented in this study is suggestive of MKs playing a role in the pathophysiology of sepsis and sepsis-related complications, larger studies are required. One particular issue to be considered when studying the MKs in blood, is the requirement for special staining to identify and quantify MKs and the failure of automated hematology analyzers to identify the MKs^68–70^ in blood Like lymphocytes, the average circulating MK diameter is between 10-15 μm in diameter and they have scant cytoplasm. MKs identified in the lungs are larger in size than those in the peripheral venous circulation^72–73^. Additionally, staining with anti-CD162 antibodies is necessary for the detection of circulating large CD61^+^CD41^+^Draq5^+^ cells. This is likely due to circulation of these cells in the form of ‘clusters’ with other peripheral blood cell types, such as neutrophils^31,74–75^. Further experiments will elucidate the participation of MK phagocytosis and chromatin web formation in pathology of infections and sepsis.

## MATERIALS AND METHODS

### Cell culture

Cord blood CD34^+^ hematopoietic stem cells were purchased and cultured in StemSpan II media with the MK supplemental cytokines (StemSpan II, Stemcell Technologies, Inc.), according to the culture and differentiation protocols from Stemcell Technologies (Stemcell Technologies Inc. Cambridge, MA). A megakaryoblastic cell line (Meg-01) was purchased and cultured in RPMI with 10% FBS, according to the standard culture protocols from ATCC (American Type Culture Collection, Manassas, VA).

### Flow cytometry

Flow cytometry was performed in order to verify appropriate cellular differentiation and for the evaluation of cell surface markers. Briefly, cells were stained with antibodies at a concentration of 1:200 for 15 minutes, with the exception of CD41, which was at a concentration of 1:100. Cells were then stained with Draq5 (Thermo Fisher Scientific, Waltham, MA) at a concentration of 1:10000 for 5 minutes. Antibodies included: anti-human CD41, HLA-ABC (MHC class I), HLA-DR (MHC class II), CD162 (p-glycoprotein-1; SELPG), CD61 (GPIIIa), CD41 (GPIIb), CD34 (Biolegend, San Diego, Ca), CD66b, and CD62P (P-selectin) (BD Biosciences, San Jose, CA). Data was obtained through the Amnis ImageStreamX Mark II imaging flow cytometer and INSPIRE Software (EMD Millipore, Billerica, MA). The accompanying IDEAS Software was used to perform data analysis. Data is reported as the percent of the total cell population that stained positive for the specific marker.

### Bacterial conjugation to pHrodo for phagocytosis experiments

Bacteria, including *Escherichia coli (E. coli), Staphylococcus aureus (S. aureus), and Streptococcus pyogenes (S. pyogenes)*, were provided by the Division of Comparative Medicine at the Massachusetts Institute of Technology (Cambridge, MA). For the killed-bacteria experiments, organisms were heat killed at 75°C for 25 minutes. Bacteria were then conjugated to pHrodo succinimidyl ester dye (Thermo Fisher Scientific) according to the manufacturer’s protocols. Briefly, bacteria were pelleted from cultures by centrifugations at 5100 rpm for 10 minutes. Pellets were resuspended in PBS (pH 9.0) at a concentration of 1×10^8^ per mL. 200 µL of this suspension was then added to 5 µL of 10 mg/mL pHrodo succinimidyl ester dye and mixed thoroughly by pipetting. Bacteria were then stained for 30 mins in the dark with gentle shaking. Following staining, 1 mL of PBS (pH 8.0) was added to the solution, and bacteria were pelleted at 13,400 rpm for 3 minutes in a benchtop centrifuge. The supernatant was removed and the pellet thoroughly resuspended in Tris Buffer (pH 8.5). The bacteria were then pelleted at 13,400 rpm for 3 mins in a benchtop centrifuge, the supernatant was removed, and the bacteria were resuspended in 1 mL of PBS (pH 7.4) before they were stored at 4°C in the dark.

### Phagocytosis inhibition experiments

*Beta-hemolytic E. coli* was provided by the Division of Comparative Medicine at the Massachusetts Institute of Technology (Cambridge, MA). Bacteria were heat-killed by incubation at 98°C for 30 mins. Unlabeled *Staphylococcus aureus* (Wood strain without protein A) BioParticles were purchased from Thermo Fisher Scientific. Bacteria and BioParticles were pelleted by centrifugations at 5100 rpm for 10 minutes at 4°C in a swing bucket centrifuge. Pellets were resuspended in PBS (pH 9.0) at a concentration of 1×10^8 per mL. 200 µL of this suspension was then added to 5 µL of 10 mg/mL pHrodo Red succinimidyl ester dye +/- Alexa Fluor 488 succinimidyl ester dye (Life Technologies) and mixed thoroughly by pipetting. Bacteria were then stained for 30 mins in the dark with gentle shaking. Following staining, 1mL of PBS (pH 8.0) was added to the solution, and bacteria were pelleted at 13,400 rpm for 3 mins in a benchtop centrifuge. The supernatant was removed and the pellet was thoroughly resuspended in Tris Buffer (pH 8.5). Again, the bacteria were pelleted at 13,400 rpm for 3 mins in a benchtop centrifuge, the supernatant was removed, and the bacteria resuspended in 1 mL of PBS (pH 7.4).

Blood was collected in ACD Vacutainer tubes (BD, Becton, Dickinson and Company, Franklin Lakes, NJ). Neutrophils were isolated from whole blood using a negative-selection protocol. Briefly, neutrophils were isolated using a density gradient with HetaSep (STEMCELL Technologies Inc. Vancouver, Canada) and then purified with EasySep Human Neutrophil Kit (STEMCELL Technologies Inc. Vancouver, Canada), following manufacturers protocol. Neutrophil purity was assessed to be >98% and cell count was performed using a hemocytometer. Neutrophils were subsequently re-suspended in the same media as the Meg-01 cells (RPMI + 10% FBS).

Meg01 and isolated neutrophils were incubated with 10 µg/mL Cytochalasin B from Dreschslera dematodia (C6762; Sigma-Aldrich) or DMSO for 30 mins prior to addition of bacteria. Cells were incubated for 2 hours with heat-killed bacteria co-labelled with pHrodo Red and Alex Fluor 488 to measure phagocytosis. 10 µL of cells were then imaged on a disposable C-Chip hemocytometer (In Cyto, SKC, Inc. C-Chip) using a 10X and 20X objective on a Nikon TiE fluorescent microscope. Stitched 6 x 6 field of view large images were de-identified and 100 cells scored blind for red fluorescence for the positive indication of pHrodo internalization and acidification. The following conditions were included in the experimental design and analysis: cells (negative control), cells with unstained bacteria (control), cells with CCB (control), cells with stained bacteria (experimental), cells with stained bacteria and CCB (experimental). No nucleated cells were noted to have a red fluorescent cytoplasm when not co-incubated with the stained bacteria. There was also no increase in the number of nucleated cells with red cytoplasm in the conditions only treated with CCB.

### Immunofluorescence Imaging

Cells were co-incubated with either the pHrodo-conjugated bacteria, Zymosan A *S. cerevisiae* BioParticles (ThermoFisher Scientific), for 60 minutes or overnight at 37°C on poly-lysine coated slides (Sigma Aldrich, St Louis, MO, USA). The slides were then rinsed gently three times with PBS and the adhered cells were subsequently stained for live imaging or fixed and then stained. For evaluation of live cells, cells were stained with Hoechst (Thermo Fisher Scientific) at a concentration of 1:2000 for 5 minutes. For the calcein green-stained cells, cells were also stained with calcein (Thermo Fisher Scientific) at 1:1000 for 5 minutes. For chromatin web evaluation, cells were also stained with SYTOX orange (Thermo Fisher Scientific) at 1:50 for 5 minutes. For histone staining of extracellular contents, cells co-incubated with *E. coli* LPS for 60 minutes at 37°C. They were then fixed with 4% paraformaldehyde (Santa Cruz Biotechnology, Dallas, TX, USA) for 30 minutes and then concentrated onto a poly-lysine coated slide using the Cytospin 4 cytocentrifuge (Thermo Fisher Scientific), for 5 minutes at 1250 rpm, rinsed once with di-water and stored at −80°C until staining and evaluation. For staining, the slides were thawed at room temperature and blocked with 5% donkey serum (Jackson Immunoresearch) for 2 hours. The slides were rinsed three times with PBS and then treated with the following primary antibodies for three hours: mouse anti-human neutrophil elastase (NE; ELA2) antibody (950334, Novus Biologicals, Littleton, CO, USA) at 1:300, rat anti-Histone H3 (phospho S28) antibody (HTA28, Abcam, Cambridge, MA, USA) at 1:500, and rabbit anti-human myeloperoxidase (A039829-2, Dako, USA) at 1:300. The slides were then rinsed three times with PBS and incubated with secondary antibodies, including donkey anti-rabbit 488, donkey anti-rat 647, and donkey anti-mouse 568 (Life Technologies) at 1:500 for 30 minutes. Slides were then rinsed three times with PBS and then covered with Vectashield antifade mounting medium with DAPI (Vector Labs, Burlingame, Ca, USA). The cells were then images with one of two fluorescent microscopes: Life Technologies EVOS FL (Thermo Fisher Scientific) or Nikon Eclipse 90i microscope (Nikon Instruments Inc., Melville, NY). Composites and videos were made and images were analyzed using either Fiji or GIMP software.

### Chromatin web release

Meg-01 and CB MK cells underwent various treatments to induce the formation of chromatin webs from MK cells. Meg-01 cells were co-incubated with various pathogenic stimuli, including *E. coli* LPS and live pHrodo conjugated *E. coli*, for 30-60 minutes at 37oc on a polylysine-coated slide. The poly-lysine slide was outlined with a hydrophobic pen prior to the experiment to set a boundary for the liquid. The slide was then rinsed three times in PBS. The slides were stained with 1:1000 Hoechst and 1:500 SYTOX orange and cover slipped. For the GFP-H2B experiments, Meg-01 cells were transfected with CellLight Histone 2B-GFP (Bacmam 2.0, ThermoFisher Scientific) for 48-72 hours and then labeled with MitoSox Red mitochondrial superoxide indicator (ThermoFisher Scientific) and Hoechst. Unstained slides were stored and evaluated with immunofluorescence techniques, as described above. The cells were imaged using one of two fluorescent microscopes: Life Technologies EVOS FL (Thermo Fisher Scientific) or Nikon Eclipse 90i microscope (Nikon Instruments Inc., Melville, NY).

### Double stranded-DNA (ds-DNA) quantification

Meg-01 cells, at 3 x 10^5 per mL, were co-incubated with various concentrations of *E. coli* LPS (20 pg/mL to 2 ug/mL) for 30 minutes at 37°C. The cells were then pelleted down at 1900 g for 10 minutes and the supernatant was immediately stored at −80°c. The experiment was performed in biological triplicates (cells from 3 different culture flasks) and in technical triplicates, on two different days. Double stranded DNA (dsDNA) was quantified using the Quant-iT™ PicoGreen™ dsDNA Assay Kit (Thermo Fisher Scientific) following the recommended protocol. Briefly, the supernatant samples were thawed and 50 uL was placed in a 96-well plate, followed by 50 uL of the aqueous working solution. A standard curve was created for a reference of extracellular chromatin. The plate was incubated at room temperature for 5 minutes and then read at standard fluorescein wavelengths (excitation ∼480 nm, emission ∼520 nm) on a SpectraMax Gemini XS plate reader (Molecular Devices, Sunnyvale, CA, USA). The mean fluorescent intensities (MFIs) were then converted to free dsDNA concentration according to the standard curve.

### Transmission Electron Microscopy

Meg-01 cells were co-incubated with live bacteria for 1 hour at 37°C. The cells were then pelleted down at 1900 g for 10 minutes. Immediately after removal of the culture medium, KII fixative (2.5% glutaraldehyde, 2.0% paraformaldehyde, 0.025 % Calcium Chloride in a 0.1M Sodium Cacodylate buffer, pH 7.4) was added to the cell/bacteria pellet, mixed, and allowed to fix for 20 minutes. The fixed sample was then prepared for both transmission electron microscopy and thin-section light microscopy and were subsequently imaged. Briefly, a rubber tipped cell scraper was used to gently remove the fixed monolayer from the plastic substrate. The samples were centrifuged, the fixative removed, replaced with buffer, and stored at 4°C until further processing. To make a cell block, the material was centrifuged again and resuspended in warm 2% agar in a warm water bath to keep the agar fluid. The material was then centrifuged again and the agar allowed to gel in an ice water bath. The tissue containing tip of the centrifuge tube was cut off resulting in an agar block with the material embedded within it. This agar block was then processed routinely for electron microscopy in a Leica Lynx™ automatic tissue processor. Subsequent processing was done using a Leica Lynx™ automatic tissue processor. Briefly, they were post-fixed in osmium tetroxide, stained En Bloc with uranyl acetate, dehydrated in graded ethanol solutions, infiltrated with propylene oxide/Epon mixtures, embedded in pure Epon, and polymerized overnight at 60°C. One-micron sections were cut, stained with toluidine blue, and examined by light microscopy. Representative areas were chosen for electron microscopic study and the Epon blocks were trimmed accordingly. Thin sections were cut with an LKB 8801 ultramicrotome and diamond knife, stained with lead citrate, and examined in a FEI Morgagni transmission electron microscope. Images were captured with an AMT (Advanced Microscopy Techniques) 2K digital camera.

### Transwell migration assay

Growth factor reduced (GFR) matrigel coated transwell inserts with 8 um pores (Biocoat Matrigel Invasion Chambers, Corning, New Jersey, USA) were thawed for 30 minutes at 37°C. 1 million Meg-01 cells were loaded in a total volume of 200 µL of RPMI with 10% FBS in the top chamber, while the bottom chamber was loaded with 600 µL of various conditions, including: RPMI with 10% FBS (negative control), 200 ng/mL SDF1-α (CXCL12, Peprotech, New Jersey, USA) (positive control), 220 pg/mL and 2.2 ng/mL *E. coli* LPS, Zymosan particles alone, and zymosan particles with 220 ng/mL *E. coli* LPS or with 200 ng/mL SDF1-α. The wells were incubated for 24 hrs. at 37°C. The cells in that migrated through the transwell insert into the bottom chamber were counted using Cellometer Vision automated cell counter (Niexcelom, Bioscience LLC., Lawrence, MA, USA).

### Microfluidic device fabrication

The microfluidic devices were manufactured using standard microfabrication techniques. The microfluidic device was designed to allow the formation of a chemical gradient in two steps, as previously described (Caroline JN, 2014). Briefly, a two-layer photoresist design (SU8, Microchem, Newton, MA), with a first and second layer that were 10.5 and 50 μm thick, were patterned on one silicon wafer via sequential photolithography masks and processing cycles according to the manufacturer’s protocols. The resulting patterned wafer was then used as a mold to produce PDMS (Polymidemethylsiloxane, Fished Scientific, Fair Lawn, NJ) devices, which were subsequently irreversibly bonded to glass slides (1×3 inches, Fisher). First, an array of circular wells (200 µm diameter, 57 µm height), connected to a side channel (10 µm width, 10.5 µm height) by orthogonal side-combs (4.5 µm width, 10.5 µm height) were primed with the following conditions: RPMI with 10% FBS (negative control), 200 ng/mL SDF1-α (CXCL12, Peprotech, New Jersey, USA) (positive control), 22 pg/mL, 220 pg/mL and 2.2 ng/mL *E. coli* LPS, zymosan particles, zymosan particles (1 million/mL) with 220 ng/mL *E. coli* LPS and zymosan particles (1 million/mL) with 200 ng/mL SDF1-α. 200 µL RPMI +10% FBS was used to wash the main channel. The diffusion of the chemoattractant from the circular wells, serving as sources, to the central channel, serving as the sink, produced the guiding gradient for the cells in the central channels. After the devices were primed and loaded, the chip was placed under vacuum for 10 minutes. Cells were then stained, loaded, and imaged for 18 hrs. every 10 minutes using time-lapse imaging on a fully automated Nikon TiE microscope with the biochamber at 37°C and 80% humidity, and in the presence of 5% carbon dioxide gas. Images were acquired automatically from distinct locations on each microfluidic device, with each image including a minimum of 3 circular wells. A minimum of 18 wells per condition were analyzed. Fiji manual tracking software (NIH) was used for the analysis of MK and platelet migration and behavior.

### MK quantification in patient blood samples

Venous blood from patients diagnosed with sepsis was collected and evaluated for circulating MKs (IRB protocol numbers, MGH No: 2014P002087; MIT No:150100681R001). A patient was categorized as septic when one of the diagnoses for the patient was ‘sepsis’ and when there was a confirmed infection, which could consist of bacterial infection, fungal infection, viral infection, or a combination thereof. Control samples were healthy donors. ‘Complicated’ sepsis was defined as the clinician-diagnosed development of acute kidney injury (AKI) or acute respiratory distress syndrome (ARDS) as documented in the patient’s medical records. Twenty-one samples were evaluated with 13 sepsis samples (age 34-75 yrs., 4 females, 9 males) and 5 control samples (age 25-50 yrs., 3 females, 1 male) (Table S1 and S2). Three of the sepsis samples (1 female and 2 males) had follow-up blood evaluation 3 days after initial blood analysis.

Cell surface markers were selected to determine cell type by differential marker expression. CD162 antibody was utilized to break up any platelet-leukocyte aggregates prior to analyzing the samples.^31^ White blood cell concentration was used as an internal control for the quantification method and whole blood spiked with MKs was used to validate surface marker identification of MKs in samples (Figure S1). Cell surface markers were selected to determine cell type by differential marker expression: CD41 (glycoprotein IIb; GPIIb) and CD61 (glycoprotein IIIb; GPIIIb) are found on the cell surface of platelets and MKs, where they form a GPIIb/IIIa complex and bind to fibrinogen and von Willebrand Factor (vWF) during platelet activation. CD45 (protein tyrosine phosphatase, receptor type, C; PTPRC; leukocyte common antigen; LCA) and CD162 (P-selectin glycoprotein ligand-1; PSGL1; SELPLG) are both present on leukocytes where CD45 hi and lo populations can be used to identify both neutrophils and lymphocytes. Draq5 is a nuclear marker that was used to differentiate between CD41^+^CD61^+^ anuclear platelets and nucleated MKs.

First, white blood cells were identified and quantified and were then compared to the total white blood cell count in the complete blood cell count (CBC) that was performed on the same blood sample at the Massachusetts General Hospital clinical pathology lab as part of the patient’s routine diagnostics in order to verify our concentration calculation methods (Figure S1A-C). Once white blood cell count was verified, cellular events staining positive for MK markers were then collected and quantified. The concentrations of MKs and leukocytes in the patient samples were then back-calculated, taking into account the total volume of sample analyzed and the initial 1:200 dilution of the blood.

In order to quantify circulating MKs in peripheral venous samples, we performed quantitative imaging flow cytometry on whole blood. In this set of experiments, we used the quantification of leukocytes in the blood as an internal methods control, with the quantity of leukocytes being compared to the automated CBC analyzer total white blood cell count to validate the cell concentration calculations. Blood collected in EDTA vacutainer tubes was diluted 1:200 in calcium-free hepes-tyrode buffer (Boston Scientific, Boston, MA, USA) with 20% volume of acid citrate dextrose (ACD, Boston Scientific). The diluted blood was then stained with CD41 PacBlue, CD61 FITC, CD45 CY5/594, and CD162 PE at 1:100-1:200 for 20 minute, and 1:1000 Draq5 for 5 minutes. The samples were then run using the Amnis flow cytometer and the data was analyzed with IDEAS software. MKs were defined as CD41^+^CD61^+^Draq5^+^ cells. Leukocytes were defined as CD162^+^CD45^+^Draq5^+^ cells. The concentrations of MKs and leukocytes in the patient samples were then back-calculated, taking into account the total volume of sample analyzed by Amnis and the initial 1:200 dilution of the blood (Equation 1).

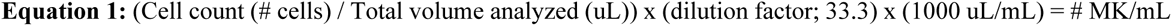

In order to explore the ability of the automated CBC analyzer to count and identify MKs, venous blood collected in EDTA was spiked with various concentrations of Meg-01 cells or cord-blood derived MKs (day 14 of differentiation). Pure MKs and the spiked whole blood samples were analyzed with flow cytometry, as described above and also run on the automated CBC analyzer in order to determine whether an automated analyzer is able to detect the presence of MKs and to see what type of cell they are categorizes as. Pure MKs were counted manually with a hemocytometer to compare with the automated analyzer (Figure S1D-E). While the imaging flow cytometer was able to specifically identify cells as MKs, the automated CBC analyzer was unable to identify them.

### Pathology samples

Histopathology samples from patients that underwent autopsies were retrospectively collected and evaluated. All samples and patient information were collected and handled according to MGH and Massachusetts Institute of Technology (Cambridge, MA, USA) IRB protocol (MGH No: 2014P002087; MIT No:150100681R001). A patient was categorized as septic when the cause of death was determined to be sepsis by the official pathology report. Control patients were defined as having primary cardiac disease as the cause of death. Fifteen samples were evaluated with 9 sepsis samples (age 60-90 yrs., 2 females, 3 males) and 5 control samples (age 68-87 yrs., 7 females, 2 males) (Table S3). All histopathology slides were de-identified and analyzed blindly.

Paraffin embedded tissue samples, including kidney, and the right middle lung lobe were sectioned and stained with either Hematoxylin & Eosin (H&E), gram stain, or with HRP-labeled CD61 antibodies by the Division of Comparative Medicine (MIT, Cambridge, MA, USA) and the Massachusetts General Hospital (Boston, MA, USA), respectively. The percent of CD61 staining per renal glomerulus was quantified using ImageJ software (NIH). Twenty renal glomeruli were evaluated for each patient. For evaluation of the lungs, the number of MKs were counted in ten 40x magnification views of the right middle lung lobe for each patient.

### Statistical Analysis

Statistics were performed using both Microsoft Excel and GraphPad Prism Software (GraphPad Software, Inc.). Either one-way ANOVA or student t-tests were performed to compare between conditions. A p-value of <0.05 was considered significant.

## Acknowledgements

The authors would like to thank Carolyn Madden at the Division of Comparative Medicine (MIT) for her isolation and preparation of the bacterial cultures used in this paper.

## Authorship Contributions

G.H.F developed the hypothesis, designed and performed experiments, and wrote the manuscript. F.E provided guidance on bacterial phagocytosis experiments and chemotaxis assays, performed phagocytosis assays and contributed to manuscript preparation. L.Z. provided guidance and pathology review of the sepsis samples, contributed to manuscript preparation. M.S. Performed electron microscopy, contributed to manuscript preparation. J.J. and A.L.M maintained cell cultures and performed chemotaxis and phagocytosis experiments and data analysis. K.W. provided guidance on immunofluorescence experiments and contributed to manuscript preparation. D.O. performed automated image analysis on the kidney pathology samples. C.V., D.I., J.G.F., and R.G.T designed experiments, analyzed the findings, and prepared the manuscript.

